# Breathe in, breathe out: Bacterial density determines collective migration in aerotaxis

**DOI:** 10.1101/2025.04.02.646741

**Authors:** Dipanjan Ghosh, Brato Chakrabarti, Xiang Cheng

## Abstract

Bacteria navigate their environment by biasing their swimming direction toward beneficial chemicals and away from harmful ones. Out of all the chemicals bacteria respond to, oxygen stands out due to its ubiquitous presence, distinct influence on bacterial metabolism and motility, and historical role in chemotaxis research. However, a coherent understanding of bacterial motility in oxygen gradients, known as *aerotaxis*, remains elusive, as evidenced by conflicting reports on the migration direction of the model organism *Escherichia coli* in self-generated oxygen gradients. Here, by combining experiments, simulations, and theory, we provide a unified framework elucidating the fundamental biophysical principle governing bacterial aerotaxis. We track the migration of bacteria in a capillary channel under self-generated oxygen gradients and show that the migration direction depends on the overall bacterial density. At high densities, bacteria migrate toward regions of higher oxygen concentration, whereas at low densities, they move in the opposite direction. We identify a critical bacterial density at which collective migration ceases, despite the presence of oxygen gradients. A kinetic theory, based on the assumption that bacteria seek an optimal oxygen concentration, is then developed to quantitatively explain our experimental findings. We validate this hypothesis by demonstrating the biased movement of individual bacteria in a dense suspension and proposing a signaling pathway that enables this behavior. Thus, by bridging the molecular level understanding of the signaling pathway, the motility of single bacteria in oxygen gradients, and the collective population dynamics shaped by oxygen diffusion and consumption, our study provides a comprehensive understanding of aerotaxis, addressing the long-standing controversy over how bacteria response to non-uniform oxygen distributions pervasive in microbial habitats.

## I. INTRODUCTION

Chemotaxis—the directed motion of microorganisms in response to chemical gradients—is a fundamental microbiological process that enables the migration of bacteria toward nutrients and away from toxins, essential for their survival, growth, and proliferation [1–3]. Although Leeuwenhoek already observed the motility of microorganisms in 1670s using his primitive single-lens microscopes, bacterial chemotaxis was not discovered until the late nineteenth century by Engelmann and Pfeffer, who first documented bacterial responses to oxygen [4–6] (for reviews of early chemotaxis research, see [7] and [8]). Subsequent studies have extensively characterized the chemotactic response of bacteria to chemoattractants such as amino acids [9], sugars [10], and dipeptides [11] and chemorepellants such as alcohols [12] and heavy metal cations [13, 14]. Despite oxygen’s historical significance in the discovery of chemotaxis, it elicits unique and complex behavior in the response of bacteria—referred to as *aerotaxis*—unobserved with other chemical stimuli.

While anaerobic bacteria stop producing ATP and cease swimming when in contact with oxygen [15], facultative anaerobes and obligate aerobes such as *Escherichia coli* (*E. coli*) and *Bacillus subtilis* (*B. subtilis*) perform aerobic respiration in the presence of oxygen that facilitates the ATP production and thereby promotes their metabolic activity [16]. Beyond the model organisms *E. coli* and *B. subtilis*, aerotactic response has been widely studied in diverse microbial species such as *Azospirillum brasilense* [17–21], *Caulobacter crescentus* [22], *Burkholderia contaminans* [23], *Shewanella oneidensis* [24–26]. The aerotaxis of oxygen-utilizing bacteria is crucial for their adaptation to environmental changes with heterogeneous oxygen distributions [27, 28]. The process has substantial medical and ecological ramification, ranging from pathogen spreading to biogeochemical nutrient cycling [28–33]. Aerotaxis can also be harnessed to optimize microbial activity in biotechnological applications such as waste treatment, fermentation, and toxicity detection [34]. Furthermore, unraveling the mechanisms of aerotaxis is critical for elucidating the emergent collective dynamics of bacteria driven by oxygen diffusion and consumption—phenomena that have gained a resurgence of research interest in recent years in the context of active matter [35–39].

Despite the great importance of bacterial aerotaxis, our understanding of this fundamental process is still incomplete. Experiments on even the model organism, *E. coli*, have yielded contradictory results regarding their response to self-generated oxygen gradients. Seminal experiments by Adler [8] and subsequent studies by Holz and Chen [40] demonstrated that a suspension of *E. coli* in a capillary tube formed a traveling band that moved toward the air-water interface. The observation led to the suggestion that bacteria are attracted to the highest available oxygen concentrations, a hypothesis that has been widely adopted to explain various collective dynamics of oxygen-utilizing bacteria [36, 41–44]. Rebbapragada and co-workers also observed a band of *E. coli* forming within a capillary channel. However, they found that this band appeared at a specific distance from the oxygen-rich air-water interface [45]. Based on this finding, they proposed that bacteria position themselves at an optimal oxygen concentration rather than at the highest available concentration. The temporal variation of the band was not reported in their study. In contrast to the two findings above, the more recent study by Douarche *et al*. found a band of *E. coli* migrating *away* from the air-water interface toward regions of low oxygen concentrations [46]. The authors suggested that this unusual phenomenon arises from changes in the diffusive behavior of bacteria at different oxygen levels. How could the same chemical—oxygen—trigger completely opposite migratory responses in a population of *E. coli* ?

In this work, we provide a unified framework on bacterial aerotaxis that resolves the long-standing controversy over the migration of *E. coli* in self-generated oxygen gradients. We systematically study bacterial migration in a capillary channel at varying bacterial densities. Our benchmark experiments show that bacteria spontaneously concentrate into a band in the channel, which, depending on the overall density of the bacterial population, moves either toward or away from the oxygen-rich interface. This unexpected finding reconciles the conflicting reports in the literature. To elucidate the biophysical mechanism underlying the opposite bacterial migration trends, we develop a kinetic theory for the bacterial population under the assumption that *E. coli* seeks an optimal oxygen concentration. With biological parameters relevant to our experiments, the theory quantitatively captures our key findings. Through an analysis of a simplified version of the model, we further demonstrate that the opposite migration trends stem from the separation of time scales between oxygen diffusion and consumption and the aerotactic movement of bacteria. Finally, we validate the assumption of optimal oxygen seeking— the cornerstone of our model—through direct visualization of the biased swimming of individual bacteria in a population and through consideration of the biochemical molecular signaling pathway supporting such a behavior. Taken together, by integrating biophysics across three distinct scales—molecular, single-bacterium, and population—we offer a comprehensive understanding of the collective migration of oxygen-utilizing bacteria in self-generated oxygen gradients. Our study provides crucial insights into bacterial motility in complex environments and lays the groundwork for understanding diverse collective dynamics in bacterial suspensions.

The paper is organized as follows: we first present our experimental findings in Sec. II and describe the kinetic theory in Sec. III. A simplified model is then constructed in Sec. IV, illustrating the competing physical factors at play. In Sec. V, we show the trajectories of individual bacteria in a suspension, offering unambiguous evidence of the optimal oxygen seeking at the single-bacterium level. Finally, we propose a signaling pathway for optimal oxygen seeking in Sec. VI, which bridges the scales between molecular signaling, the motility of a single bacterium, and the collective population dynamics, thus providing a comprehensive description of bacterial aerotaxis. The conclusion and future work are discussed at the end.

## II. EXPERIMENTS

### A. Setup and results

Experiments on bacterial chemotaxis have focused either on the motion of single cells in dilute suspensions [47–50] or on the collective migration of bacterial populations [51–58]. The chemotactic behavior of individual bacteria within a dense population remains largely unexplored. Here, we examine simultaneously the response of bacteria to oxygen gradients at both the single-cell and population levels by introducing a subpopulation of fluorescently tagged *E. coli* in a bacterial suspension of controlled density. Apart from the fluorescent plasmid insertion, the genotypes of the fluorescent and non-fluorescent cells are identical, ensuring that both types of cells have identical responses toward oxygen. Bacterial density is kept low enough to preclude the transition to collective bacterial swimming [59, 60].

Figure 1(a) illustrates our experimental setup, where a suspension of mixed *E. coli* is introduced in a horizontal glass capillary channel with one end open to the atmosphere at *x* = 0. Oxygen diffuses from the air-water interface into the channel, which is continuously consumed by bacteria, creating an oxygen gradient with the highest concentration at the interface. At the population level, we determine the spatial density profile of bacteria, *ρ*(*x*), by counting fluorescently tagged cells at each position across the channel (Fig. 1(b)). *ρ*(*x*) is normalized by the total number of detected bacteria in each sample to eliminate the influence of variations in the ratio of fluorescent to non-fluorescent bacteria. Thanks to the relatively low concentration of fluorescent cells in the background of densely populated, unlabeled bacteria, we are able to track the dynamics of individual bacteria. Details on our bacterial culture, experimental setup, and image analysis can be found in Appendix A-D.

**FIG. 1.**
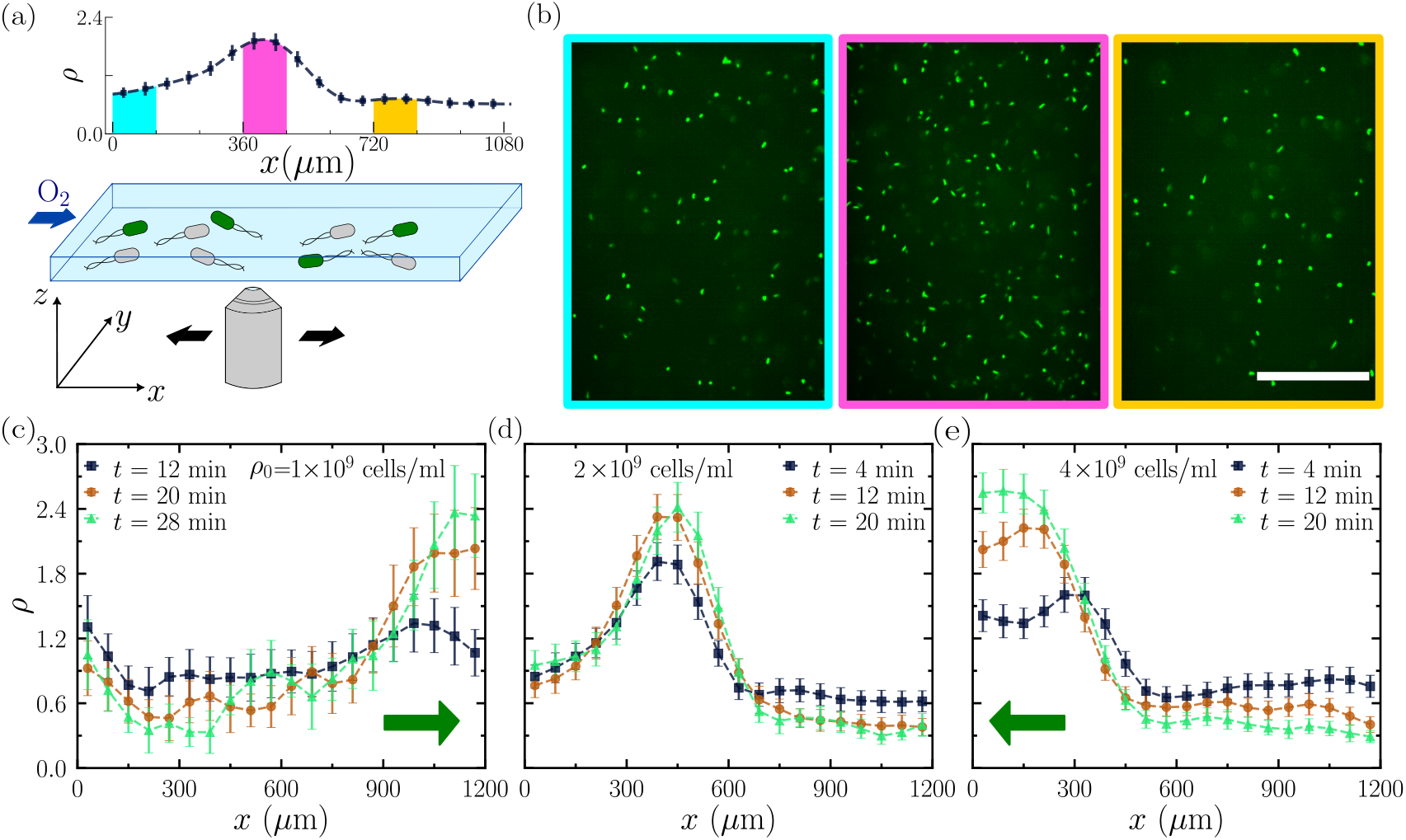
Bacterial aerotaxis in experiments. (a) Schematic of the experimental setup with a suspension of *E. coli* in a glass capillary channel in contact with atmospheric oxygen at *x* = 0. The objective lens moves along the *x* direction to record videos at different spatial locations. Top: Spatial bacterial density profile from an experiment with overall cell density *ρ*_0_ = 2 × 10^9^ cells/ml at *t* = 4 minutes. (b) Panels outlined with different colors show snapshots of the suspension at *x* = 0 − 120 *µ*m, 360 − 480 *µ*m, and 720 − 840 *µ*m, corresponding to the shaded regions on the density profile in (a). Fluorescently tagged bacteria, imaged using confocal microscopy, appear as bright ellipsoids against a background of non-fluorescent bacteria. The ratio of fluorescent to non-fluorescent cells is 1:4. The scale bar is 50 *µ*m. The temporal variation of bacterial density profiles for *ρ*_0_ = 1 × 10^9^ cells/ml (c), 2 × 10^9^ cells/ml (d), and 4 × 10^9^ cells/ml (e). The migration directions of the bacterial population in (c) and (e) are shown using the green arrows. Error bars indicate the standard error. *ρ*_0_ = 4 × 10^9^ cells/ml (Fig. 1(e)), the peak migrates in the opposite direction toward the air-water interface, where oxygen concentration is highest. At an intermediate density *ρ*_0_ = 2 × 10^9^ cells/ml (Fig. 1(d)), the peak position remains nearly stationary.

We vary the overall density of cells, *ρ*_0_, and measure the temporal variation of the spatial distribution of bacteria *ρ*(*x*). Figure 1(c)-(e) show *ρ*(*x*) at different times for three different *ρ*_0_. There are two interesting observations. First, regardless of *ρ*_0_, we observe the emergence of a high-density peak in *ρ*(*x*), i.e., a bacterial band, at early times. The peak appears closer to the interface at larger *ρ*_0_. Second, the peak migrates over time, with its direction of movement depending on *ρ*_0_. At a low density *ρ*_0_ = 1 × 10^9^ cells/ml (Fig. 1(c)), bacteria migrate away from the air-water interface toward regions of low oxygen concentrations. In contrast, at a high density

To quantify different migration behaviors, we measure the average migration speed of the bacterial population *v*_*m*_ as a function of *ρ*_0_ (Fig. 2 and Appendix E). *v*_*m*_(*ρ*_0_) shows a clear transition from interface-bound to bulk-bound migration at a critical density of *ρ*_*c*_ ≈ 1.5 ×10^9^ −2.0 × 10^9^ cells/ml. A greater deviation from *ρ*_*c*_ results in faster migration either toward or away from the interface. As we will discuss next, these results defy explanation by existing chemotaxis models and challenge the current understanding of the migration of aerotactic bacteria in self-generated oxygen gradients.

**FIG. 2.**
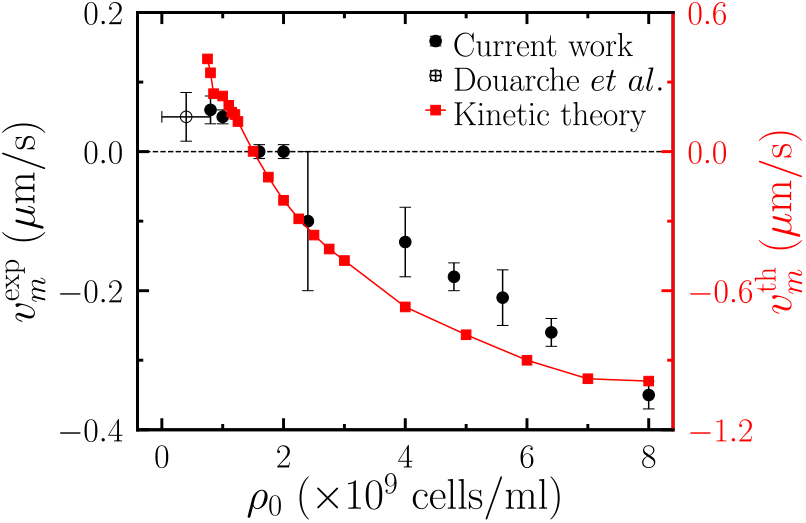
Migration speed of bacterial population from experiments 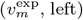 and kinetic theory 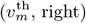 as a function of overall bacterial densities *ρ*_0_. Filled black circles represent our experimental data, while the empty circle corresponds to the measurement by Douarche *et al*. [46]. Error bars indicate the standard error.

### B. Existing theories

The formation and migration of a bacterial band are among the most prominent and well-studied features in the collective dynamics of bacteria in chemotaxis [8, 61– 70]. In the classic Keller-Segel model [66], the migration speed of bacteria in response to a spatial gradient of a chemoattractant along the *x*-axis is given as *v*_*x*_(*x*) = (*χ/c*)*∂*_*x*_*c*, where *χ >* 0 quantifies the strength of the chemotactic response and *c*(*x*) is the chemoattractant concentration. In the geometry of our experiments, consider two points *x*_1_ and *x*_2_ with 0 *< x*_1_ *< x*_2_. The oxygen concentration satisfies *c*(*x*_1_) *> c*(*x*_2_), independent of bacterial density *ρ*_0_. In a shallow gradient where *∂*_*x*_*c*(*x*_1_) ≈ *∂*_*x*_*c*(*x*_2_) *<* 0, we have *v*_*x*_(*x*_2_) *< v*_*x*_(*x*_1_) *<* 0. The above argument suggests that, while all bacteria move toward the interface, the magnitude of the migration speed of bacteria is smaller near the interface, resulting in the accumulation of bacteria and the formation of an interface-bound traveling band. Although this prediction qualitatively matches our observations for *ρ*_0_ *> ρ*_*c*_, it fails to explain the switch of migration direction for *ρ*_0_ *< ρ*_*c*_.

Douarche *et al*. proposed a chemotaxis-independent mechanism to explain the migration of bacterial bands away from the air-water interface [46]. Bacteria near the air-water interface exhibit higher motility due to high oxygen concentration, leading to a greater effective translational diffusivity compared to bacteria in the oxygen-deprived regions deep inside the channel. These authors argued that bacteria moving from high-oxygen regions slow down as they enter low-oxygen regions, causing an accumulation of bacteria and the formation of the band. As oxygen gradually diffuses into the channel, the bacterial band migrates from the interface into the bulk with a speed proportional to the difference in bacterial diffusivity between high and low-oxygen regions. Similar to the Keller-Segel model, this mechanism only solves half of the puzzle and cannot explain the interface-bound migration when *ρ*_0_ *> ρ*_*c*_. More fundamentally, the mechanism overlooks the flux created by the density gradient across the band, which counteracts the flux due to the change in effective diffusivity. It has been shown that spatial variations in diffusivity alone cannot produce non-uniform bacterial density distributions at steady state [71].

## III. KINETIC THEORY

### A. Model

To rationalize our experimental findings and explain the direction reversal of bacterial migration at *ρ*_*c*_, we develop a two-dimensional (2D) kinetic theory [36, 72– 75] to describe the dynamics of a bacterial population in a self-generated oxygen gradient. In this framework, the bacterial population is described by a coarse-grained probability density function *ψ*(**x**, *θ, t*) that characterizes the probability of finding a bacterium at **x** = {*x, y*} with orientation **p** = (cos *θ*, sin *θ*) at time *t* (Fig. 3(a)). The distribution function satisfies a conservation equation,

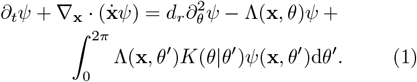

**FIG. 3.**
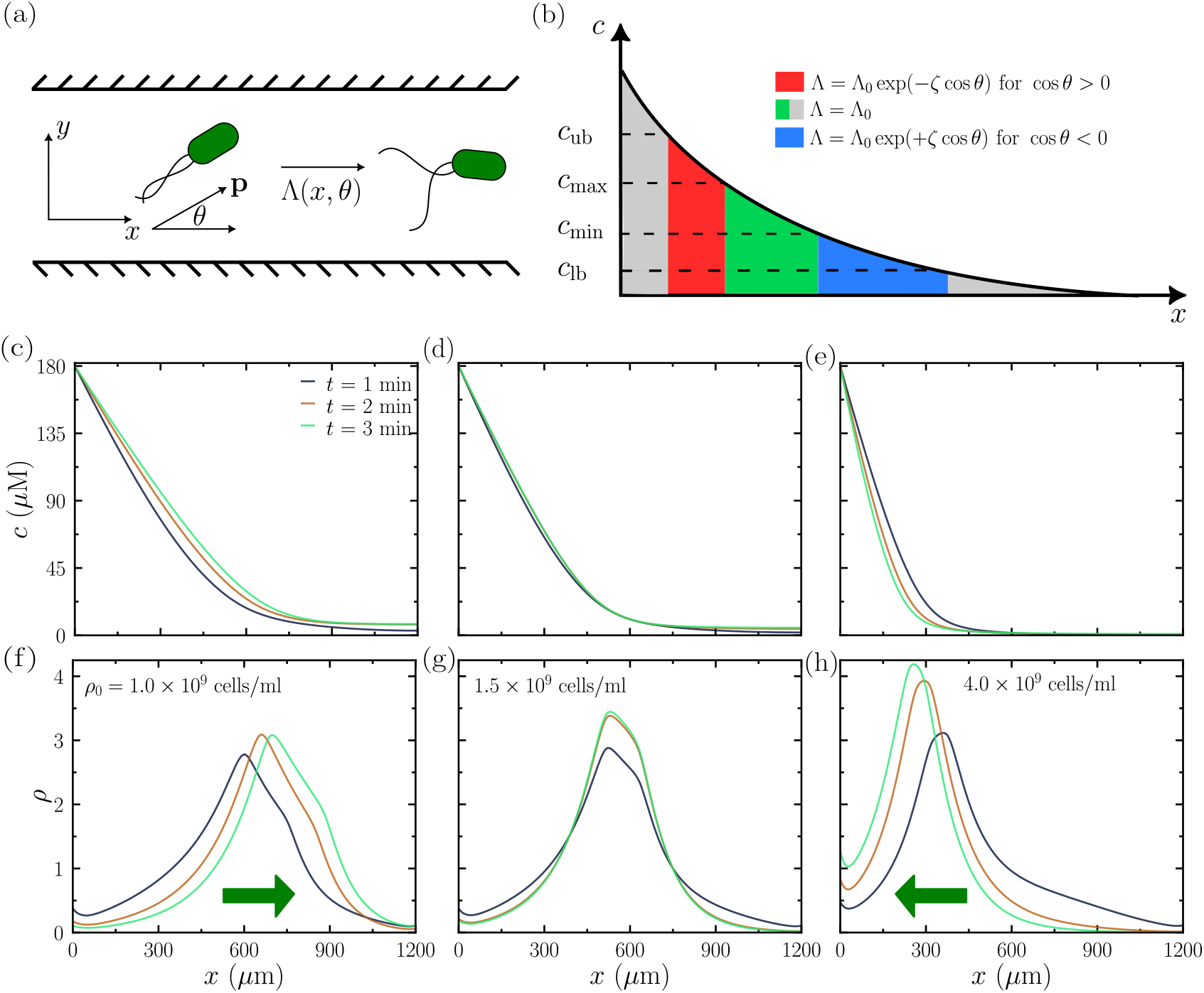
Bacterial aerotaxis as predicted by kinetic theory. (a) Schematic of the run-and-tumble of bacteria in a 2D channel. (b) Schematic illustrating variations in tumbling frequency Λ at different oxygen concentrations *c*(*x*) (not to scale). (c-e) Temporal evolution of oxygen concentration profiles and (e-g) bacterial density profiles for overall bacterial densities *ρ*_0_ = 1 × 10^9^ cells/ml (c, f), 1.5 × 10^9^ cells/ml (d, g), and 4.0 × 10^9^ cells/ml (e, h). The migration directions of the bacterial population in (f) and (h) are shown using the green arrows.

Here, 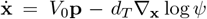 is the translational flux, where *V*_0_ is the bacterial swimming speed and *d*_*T*_ is the translational diffusivity. The first term on the right-hand side represents changes in bacterial orientation due to rotational diffusion with diffusivity *d*_*r*_. The last two terms capture the run-and-tumble dynamics of a bacterium with tumbling frequency Λ(**x**, *θ*), which depends on the instantaneous position and orientation of a bacterium. In particular, −Λ*ψ* denotes the fraction of bacteria tumbling away from orientation *θ* and the integral provides a measure of bacteria tumbling into the configuration *θ* from *θ*^′^. The transition probability *K*(*θ* |*θ*^′^) denotes the conditional probability density of tumbling from orientation *θ*^′^ to *θ* in the absence of external tumbling cues. In this work, we assume de-correlated tumbles with *K*(*θ*| *θ*^′^) = 1*/*(2*π*).

As oxygen diffuses into the channel from the air-water interface, setting up a gradient in the *x* direction, bacterial density depends on *x* alone. Neglecting variation in the *y* direction, Eq. (1) simplifies to

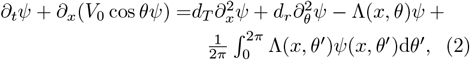

where *x* ∈ [0, *L*] with *L* denoting the length of the channel. The probability density is normalized such that 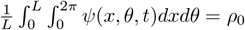. The local density of bacteria at location *x* is obtained by integrating out the orientational degree of freedom, 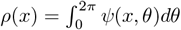. The above conservation equation is accompanied by a no-flux boundary conditions at both ends of the channel

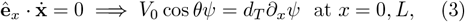

ensuring the conservation of the total number of bacteria.

In conventional chemotaxis, bacteria swim up the gradient of a chemoattractant by reducing the tumbling frequency Λ, which, as shown in Sec. II B, cannot explain the present experimental observations. Here, we adopt a different hypothesis: instead of swimming toward regions of higher oxygen concentrations, *E. coli* seek an intermediate concentration of oxygen, optimal for their internal metabolism [45, 76]. It has been proposed that *E. coli* cannot directly sense the external oxygen level in their environment. The local concentration of oxygen, *c*(*x*), affects the internal metabolic state or energy level of the bacteria. The optimal internal energy state of the bacterium is achieved for oxygen concentrations between *c*_min_ *< c < c*_max_, where *c*_min_ and *c*_max_ are the lower and upper favorable oxygen concentrations, respectively. An *E. coli* bacterium modifies its tumbling frequency to swim toward the regions of the favorable oxygen concentrations [45].

To implement this hypothesis in the kinetic theory, we extend the model proposed by Mazzag *et al*. for the 1D swimming of aerotactic bacteria *Azospirillum brasilense* [20] to the 2D run-and-tumble dynamics of *E. coli*. The tumbling frequency Λ(*x, θ*) is given as

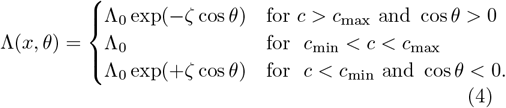

Here, Λ_0_ is the baseline tumbling frequency of bacteria in the absence of any chemical gradient and *ζ >* 0 represents the sensitivity of the tumbling frequency to the local oxygen concentration. In addition, we assume that bacteria can only sense and respond to oxygen when the oxygen concentration is less than an upper bound *c < c*_ub_ but greater than a lower bound *c > c*_lb_. Outside this regime, Λ = Λ_0_. Figure 3(b) presents a schematic depicting the variation of Λ along a typical oxygen concentration profile.

Finally, the diffusion and consumption of oxygen are given by

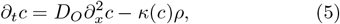

where *D*_*O*_ is the diffusivity of oxygen in water, and *κ*(*c*) is the consumption rate of oxygen by bacteria given as [20, 46, 55]

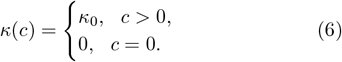

The oxygen concentration at the air-water interface is fixed at the saturation concentration *c*(0) = *c*_0_, whereas at the interior end of the channel, we prescribe a no-flux boundary condition, *∂*_*x*_*c*(*L*) = 0.

The following parameters are used in our theory. The baseline tumbling frequency is chosen as the inverse of the average run time of bacteria, Λ_0_ = 1 s^−1^. The swimming speed of bacteria is *V*_0_ = 20 *µ*m/s. The translational and rotational diffusivity of bacteria is taken to be *d*_*T*_ = 4 × 10^−10^ m^2^/s and *d*_*r*_ = 0.05 s^−1^, respectively [36]. The concentration of oxygen at the interface is given by the saturation oxygen concentration in water at room temperature *c*_0_ = 180 *µ*M [77]. We set *c*_ub_ = 0.9*c*_0_, *c*_max_ = 0.1*c*_0_, *c*_min_ = 0.05*c*_0_, and *c*_lb_ = 5 × 10^−5^*c*_0_, in close agreement with unpublished data referenced in [18] and [76]. The tumble sensitivity, *ζ*, is set to 0.8, which yields a range of bacterial run length consistent with experiments. Finally, the molecular diffusivity of oxygen is *D*_*O*_ = 2.4 × 10^3^ *µ*m^2^/s and the oxygen consumption rate is *κ*_0_ = 1.2 × 10^−9^*µ*M/[(cells/ml) · s] [55]. Notably, all parameters are chosen to be consistent with independent experimental measurements reported in prior literature. There are no free parameters in our model.

We solve Eqs. (2) and (5) using the open source spectral solver Dedalus [78] with a domain size *L* = 1200 *µ*m at different values of overall density *ρ*_0_. Following the experiments, the initial density of bacteria, *ρ* = *ρ*_0_, is spatially uniform, and the initial oxygen concentration in the channel is *c* = 0.

### B. Results

The kinetic theory reproduces the key findings of our experiments. Figure 3(c)-(e) and (f)-(h) show respectively the temporal variations of the oxygen concentration profile *c*(*x*) and bacterial density profile *ρ*(*x*) for different overall density *ρ*_0_. Independent of *ρ*_0_, oxygen diffuses from the air-water interface at *x* = 0 and is consumed by bacteria as it enters the channel, leading to a monotonically decreasing spatial profile. Meanwhile, bacteria rapidly accumulate to form a density peak. As *ρ*_0_ increases, the initial position of the bacterial density peak moves closer to the interface.

At a low *ρ*_0_ = 1 × 10^9^ cells/ml, the oxygen profile *c*(*x*) penetrates progressively deeper into the channel away from the air-water interface over time (Fig. 3(c)). Alongside this shift, the peak of the bacterial density also migrates away from the interface toward the region of low oxygen concentrations (Fig. 3(f)). At an intermediate *ρ*_0_ = 1.5 × 10^9^ cells/ml, both *c*(*x*) and *ρ*(*x*) remain stationary (Figs. 3(d) and (g)). Finally, at a higher density *ρ*_0_ = 4 × 10^9^ cells/ml, *c*(*x*) retreats toward the air-water interface over time (Fig. 3(e)), accompanied by bacterial migration toward the region of higher oxygen concentrations at the interface (Fig. 3(h)).

We extract the position of the density peak *x*_*p*_ and the oxygen decay length ℒ_*O*_ defined as the length, where *c*(*x* = ℒ_*O*_) = *e*^−1^*c*_0_. Figure 4 shows the temporal evolution of *x*_*p*_ and ℒ_*O*_ for different *ρ*_0_. At low *ρ*_0_, *x*_*p*_ increases monotonically (Fig. 4(a)), indicating a migration of the bacterial population away from the air-water interface toward the region of low oxygen concentrations. At a critical density *ρ*_*c*_ ≈1.5 × 10^9^ cells/ml, *x*_*p*_ remains constant after an initial transient period of rapid increase, indicating no directional migration. Finally, for *ρ*_0_ *> ρ*_*c*_, *x*_*p*_ increases transiently at short times and then gradually migrates toward the air-water interface into the oxygen-rich region. The trend of the oxygen decay length ℒ_*O*_ closely follows that of *x*_*p*_ (Fig. 4(b)). For *ρ*_0_ *< ρ*_*c*_, ℒ_*O*_ increases monotonically as oxygen continuously diffuses into the bulk. In contrast, for *ρ*_0_ *> ρ*_*c*_, oxygen initially penetrates into the channel, followed by a retreat at later times toward the air-water interface driven by the consumption of bacteria.

**FIG. 4.**
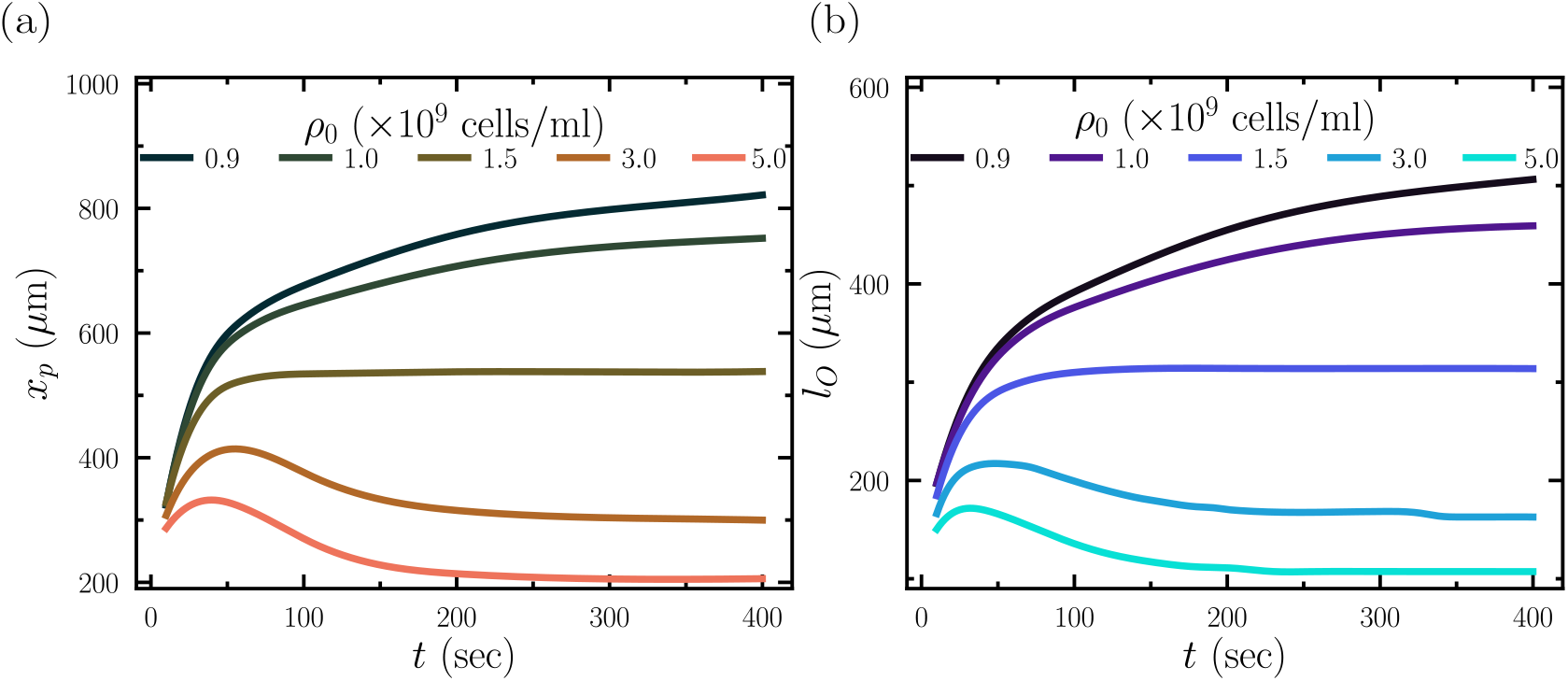
Temporal evolution of (a) bacterial density peak *x*_*p*_, and (b) oxygen decay length *l*_*O*_ from the kinetic theory at different overall cell densities *ρ*_0_.

Using the same procedure as in the experimental analysis, we compute the average bacterial migration speed from the kinetic theory, *v*_*m*_(*ρ*_0_). We exclude the initial transient period, which falls outside the temporal resolution of our experiments (Appendix E). Figure 2 compares the predictions of the kinetic theory with experimental measurements. With experimentally determined parameters, the predicted speed is within a factor of three of the experimental values. More importantly, the theory shows the correct trend of *v*_*m*_(*ρ*_0_) and accurately predicts the critical density *ρ*_*c*_.

Thus, our combined experimental and numerical findings quantitatively resolve the long-standing debate over whether *E. coli* migrates toward [8, 40] or away from the air-water interface. By identifying the overall density, *ρ*_0_, as the key control parameter, we establish a unifying framework that reconciles these seemingly conflicting observations. Furthermore, our results provide a strong support to the hypothesis that *E. coli* seeks an optimal intermediate oxygen concentration during aerotaxis. While aerotaxis naturally drives the formation of a bacterial band at the region with optimal oxygen concentrations, the direction of band migration emerges from a non-trivial interplay between oxygen diffusion and consumption and evolving non-uniform bacterial density distribution. To provide physical insights into this complex process, we construct a minimal model to illustrate the competing physical factors at play.

## IV. MINIMAL MODEL

The construction of a minimal model to explain bacterial migration in self-generated oxygen gradients hinges upon two observations from the kinetic theory: (a) fast oxygen diffusion and consumption at early times, and (b) slow aerotaxis-driven bacterial movement at later times. We discuss the physics at these two timescales.

### A. Short-time oxygen diffusion and comsumption

Our experiments and simulations reveal that the aerotactic response of bacteria occurs much more slowly than the diffusion and consumption of oxygen in the capillary channel. To illustrate this separation of time scales, we first estimate the penetration depth of oxygen *x* = ℒ_*p*_, at which oxygen runs out due to the consumption of uniformly distributed bacteria. The penetration depth is computed by solving

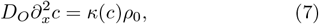

subjected to the boundary conditions *c*(0) = *c*_0_ and *∂*_*x*_*c*(*L*) = 0. By positing that the oxygen concentration falls to zero at *x* = ℒ_*p*_ ≤ *L, c*(*x*) takes the form:

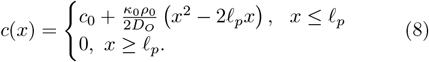

The solution yields the penetration depth

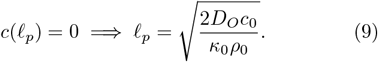

The time for oxygen to diffuse from the interface over a distance of ℒ_*p*_ is 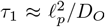. Since the optimal oxygen concentration preferred by *E. coli* is much smaller than the interfacial oxygen concentration (*c*_max_ ≪ *c*_0_), the initial bacterial density peaks around ℒ_*p*_. The time for an aerotactic bacterium to migrate from the interface to the optimal location is then *τ*_2_ ℒ_*p*_*/v*_*b*_, where *v*_*b*_ is the migration speed of aerotactic bacteria. We approximate *v*_*b*_ using the characteristic migration speed of bacterial population ∼ 0.3 *µ*m/s (Fig. 2). Using the critical bacterial density *ρ*_*c*_ = 1.5 × 10^9^ cells/ml as the density scale, we have *τ*_2_*/τ*_1_ = *D*_*O*_*/*(ℒ_*p*_*v*_*b*_) ≈ 12. Choosing a smaller *v*_*b*_ would further increase this ratio. Thus, around the critical density *ρ*_*c*_, the time scales of oxygen diffusion and consumption and bacterial migration are well separated. The location of the bacterial density peak at early times, 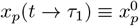, is determined by solving Eq. (8) for 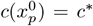, where *c*^*^ is the optimal oxygen concentration, yielding,

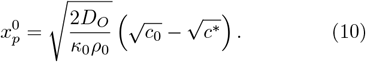

As 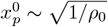, the initial position of the band is closer to the interface at higher *ρ*_0_, consistent with our experimental observation and the kinetic theory prediction (Figs. 1 and 3). To verify the prediction quantitatively, we estimate the position of the density peak at early times in our kinetic theory. Specifically, we compute *τ*_1_ for each overall density *ρ*_0_ and measure the peak position *x*_*p*_ at *τ*_1_. The results quantitatively agree with the prediction of Eq. (10) (Fig. 5), confirming the timescale separation in aerotaxis. Here, we choose *c*^*^ = (*c*_max_ + *c*_min_)*/*2 = 0.075*c*_0_. The peak position from the kinetic theory exhibits a larger deviation from the predicted value at high 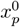, a situation corresponding to low overall density *ρ*_0_. As *ρ*_0_ decreases, ℒ_*p*_ and consequently *τ*_1_ increase, with *τ*_1_ approaching the slow bacterial migration time scale *τ*_2_. As a result, the distinction between the two time scales becomes less pronounced at low *ρ*_0_.

**FIG. 5.**
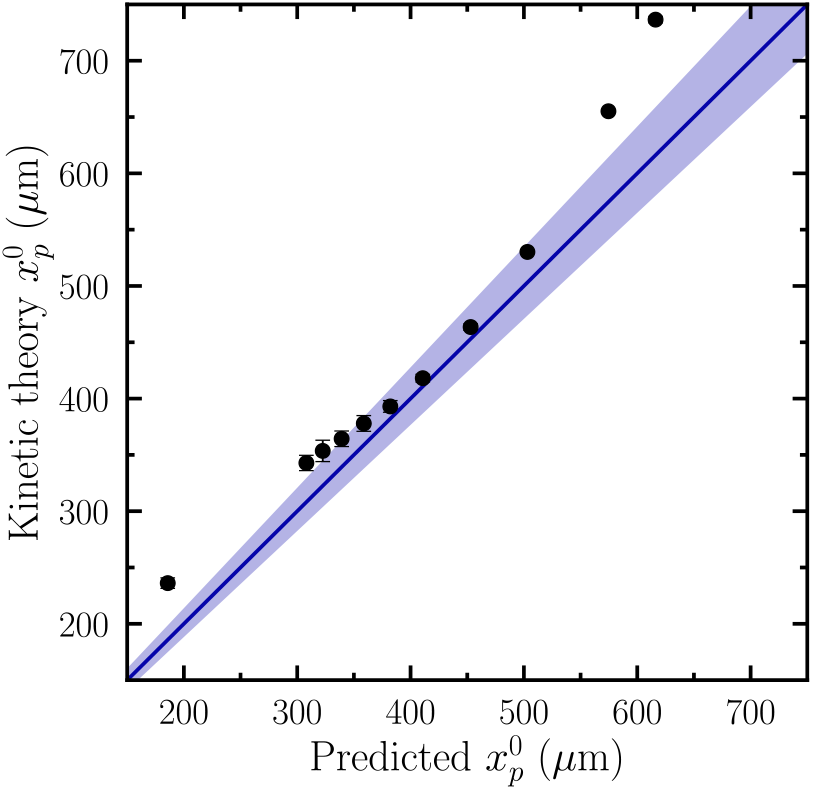
Initial peak positions 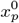 from the kinetic theory versus the predicted position from short-time oxygen diffusion and consumption (Eq. (10)). Symbols represent the average 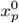 over a 20-second interval centered around *τ*_1_ from the ki-netic theory. Error bars are standard deviations. The line indicates the identity line when *c*^*^ = 0.075*c*_0_. The shaded region denotes the predicted 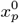 for 0.05*c*_0_ *< c*^*^ *<* 0.1*c*_0_.

### B. Long-time bacterial aerotaxis

After identifying the initial position of the density peak at 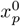, we now proceed to estimate the steady-state bacterial density profile. To this end, we consider a 1D telegrapher’s equation model for the bacteria population and define two sub-populations of bacteria moving along the +*x* and − *x* directions with probability densities *P* ^+^ and *P* ^−^, respectively. Their evolution obeys

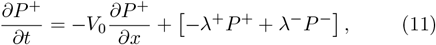

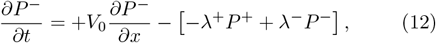

where *λ*^+^ (*λ*^−^) is the rate at which a bacterium moving along the +*x* (−*x*) direction reverses its swimming direction, capturing the run-and-tumble dynamics of *E. coli*. These transport equations are complemented by the no-flux boundary conditions *P* ^+^(0) = *P* ^−^(0) and *P* ^+^(*L*) = *P* ^−^(*L*) that ensures conservation of total number of bacteria. Adding and subtracting Eqs. (11) and (12), we obtain

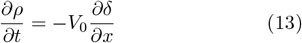

and

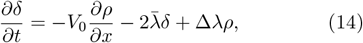

where *ρ* = *P* ^+^ + *P* ^−^ is the local density of bacteria, *δ* = *P* ^+^ − *P* ^−^ is the local density difference of the two sub-populations, 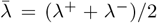 is the local average reversal frequency, and Δ*λ* = *λ*^−^ − *λ*^+^ is the mismatch between the reversal frequencies of the left- and right-going subpopulations.

Assuming that the density difference *δ* equilibrates faster compared to the density *ρ* [79–81], a quasi-static approximation of *∂*_*t*_*δ* ≈0 gives 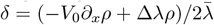 from Eq. (14). Substituting *δ* in Eq. (13) results in an advection-diffusion equation for the bacterial population

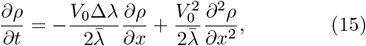

where 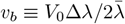 is the migration speed of aerotactic bacteria. At steady state, Equation (15) becomes

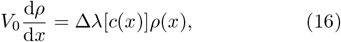

which is subjected to the conservation of bacterial number 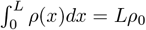.

The equation is coupled to the steady state oxygen profile *c*(*x*) determined self-consistently through

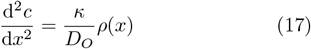

with the boundary condition, *c*(0) = *c*_0_ and *∂*_*x*_*c*(*L*) = 0, and *κ* given by Eq. (6). For a given choice of Δ*λ*(*c*), Eqs. (16) and (17) yield a nonlinear boundary value problem for the unknown density *ρ*(*x*) and oxygen profile *c*(*x*).

From Eq. (16), the steady-state density profile can be written as,

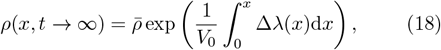

where the constant 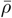 is a normalization constant. The maximum of *ρ*(*x, t* → ∞) gives the peak position of the bacterial band at steady state, 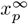.

Before solving Eq.(18), we first examine the general features of the equation, particularly how it depends on the spatial variation in bacterial tumbling dynamics relative to the optimal oxygen concentration, *c*(*x* = *x*^*^) = *c*^*^. To the left of *x*^*^, bacteria moving in the +*x* direction tumble less frequently than those moving in the −*x* direction, resulting in biased motion toward *x*^*^. To the right of *x*^*^, this trend reverses. This bias in tumbling behavior creates a mismatch between the tumbling frequencies of the two populations, quantified by Δ*λ*, as described in Ref. [20]. Specifically, Δ*λ* changes sign from positive to negative as *c* decreases past *c*^*^ with increasing *x*. Consequently, the integral 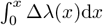 reaches a peak at *x*^*^. According to Eq.(18), *ρ*(*x*) also peaks at *x*_*p*_ = *x*^*^. Thus, the sign change of Δ*λ* at *x*_*p*_ = *x*^*^ is crucial for the formation of the bacterial band. Figure 6 illustrates the normalized Δ*λ* from kinetic theory, confirming the sign change of Δ*λ* at the peak *x*_*p*_ (Appendix F).

**FIG. 6.**
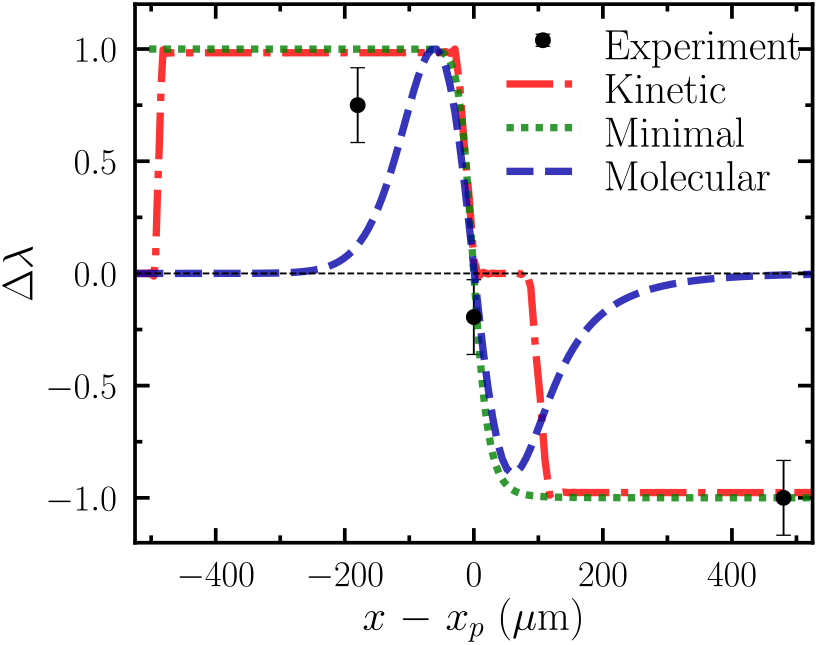
Spatial profile of tumbling frequency difference, Δ*λ*, from experiments and different models. The position of each dataset is shifted by subtracting the corresponding location of the bacterial density peak *x*_*p*_. Experimental Δ*λ* (black disks) are calculated from individual bacterial trajectories in Fig. 8. Error bars represent the standard error. All the results are normalized by their maximum absolute values. The curves from the models are calculated at *ρ*_0_ = 1.5 × 10^9^ cells/ml at their steady states. The black horizontal dashed line corresponds to Δ*λ* = 0.

### C. Migration of bacterial band

The position of the steady-state bacterial density peak, 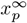, depends on the overall bacterial density *ρ*_0_. Comparing 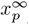 with the initial position of the density peak 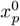 from Eq. (10) leads to three natural outcomes. (*i*) When 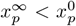, the steady-state position of the peak is to the left of its initial position, and the bacterial band migrates toward the interface over time. (*ii*) When 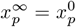, the peak remains stationary. (*iii*) When 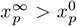, the peak moves into the bulk.

To calculate 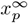 from Eqs. (16) and (17) (or equivalently Eq. (18)), we need to have an expression of Δ*λ*(*c*). Without loss of generality, we choose Δ*λ* = *α* tanh[*γ*(*c* −*c*^*^)], capturing the sign change in Δ*λ* at *c* = *c*^*^. Here, *γ* determines the steepness of the sign change of Δ*λ* near *c*^*^ and can be varied over several orders of magnitude without significantly affecting the critical bacterial density *ρ*_*c*_ (Appendix G). The amplitude of Δ*λ* is controlled by *α*, which in turn sets the advection speed of aerotactic bacteria 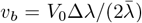 (Eq. (15)). Using *V*_0_ = 20 *µ*m/s and 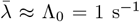 and choosing a characteristic migration speed based on experiments *v*_*b*_ ≈ 0.3 *µ*m/s (Fig. 2), we have *α* = 0.03 s^−1^. Setting *γ* = 0.33 *µ*M^−1^, we calculate 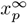 as a function of *ρ*_0_ and compare it with the peak at the early time 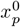. Figure 7 shows that 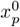 and 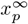 intersect at *ρ* = 1.5 × 10^9^ cells/ml, indicating a reversal in migration direction, in quantitative agreement with our experimental observation and the kinetic theory prediction on the critical bacterial density *ρ*_*c*_. Increasing *α* would enhance the advection speed of bacteria, making the migration timescale *τ*_2_ comparable to the oxygen diffusion timescale *τ*_1_. Numerical calculations show no reversal of the migration direction at large *α* within the range of *ρ*_0_ of our experiments (Appendix G). Thus, our results further confirm that, in addition to the sign change of Δ*λ* (Fig. 6), the separation of timescales is another key factor driving the observed migration dynamics.

**FIG. 7.**
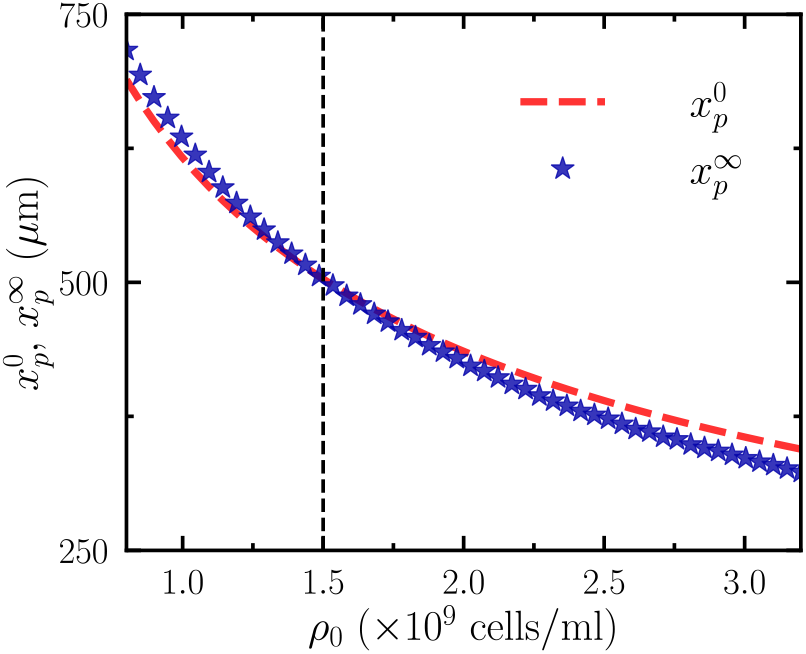
Initial peak position, 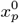, and the steady-state peak position, 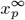, as function of mean bacterial density *ρ*_0_ from the minimal model. The vertical dashed line indicates the critical density where 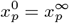.

## V. DYNAMICS OF SINGLE BACTERIA

A key assumption of our theory and model is that *E. coli* seek an optimal, intermediate oxygen concentration during aerotaxis. We validate this assumption by experimentally tracking the swimming of individual bacteria within a dense population migrating in an oxygen gradient (this section) and by proposing a molecular signaling pathway that supports this behavior (next section). In both cases, we aim to demonstrate the sign change of Δ*λ*—the difference between the tumbling frequencies of left- and right-moving bacteria—across the peak of the density profile. As shown in Sec. IV C, this sign change is the underlying mechanism driving the formation of the bacterial band during aerotaxis.

In our experiments, a bacterium enters the imaging plane of confocal microscopy through a tumble, swims within the plane at a speed *V*_0_, and exits the plane through another tumble [82]. Thus, the total distance traveled by a bacterium within the imaging plane captures its run length *R* = *V*_0_*/λ*. We measure the run length *R* and the swimming direction *θ* of bacteria and analyze the directional bias in individual bacterial runs. In particular, we focus on individuals with large *x*-velocity components, as these cells experience significant changes of oxygen concentration. Bacteria oriented with *θ* ∈ [−*π/*4, *π/*4] swim away from the air-water interface, while those with *θ* ∈ [3*π/*4, 5*π/*4] swim toward it. Here, *θ* is calculated based on the orientation of the line segment connecting the start and end points of the trajectory of a bacterium with respect to the *x* axis. We denote their respective run lengths as *R*^+^ and *R*^−^.

Figure 8 shows the distribution of *R*^+^ and *R*^−^ from over 3,300 single-cell trajectories at an early time *t* = 4 min with the overall bacterial density *ρ*_0_ = 4 × 10^9^ cells/ml. The peak of bacterial density is located at *x*_*p*_ *≈* 300 *µ*m. We categorize the trajectories based on their spatial position relative to this peak: to the left (0 *< x <* 240 *µ*m), in the middle (240 *< x <* 360 *µ*m), and to the right of the peak (360 *< x <* 1200 *µ*m). In each region, the orange and green violin plots represent the distributions of *R*^+^ and *R*^−^, respectively. Using a two-sided Kolmogorov-Smirnov test, we test the null hypothesis that the distributions of *R*^+^ and *R*^−^ are similar. On the left side of the peak, the distribution of *R*^+^ has a longer tail than that of *R*^−^, with a statistically significant difference (*p <* 0.01). In contrast, on the right side of the peak *R*^−^ shows a significantly longer tail compared to *R*^+^ (*p <* 0.01). In the middle region, the difference between the distributions of *R*^+^ and *R*^−^ is not statistically significant.

**FIG. 8.**
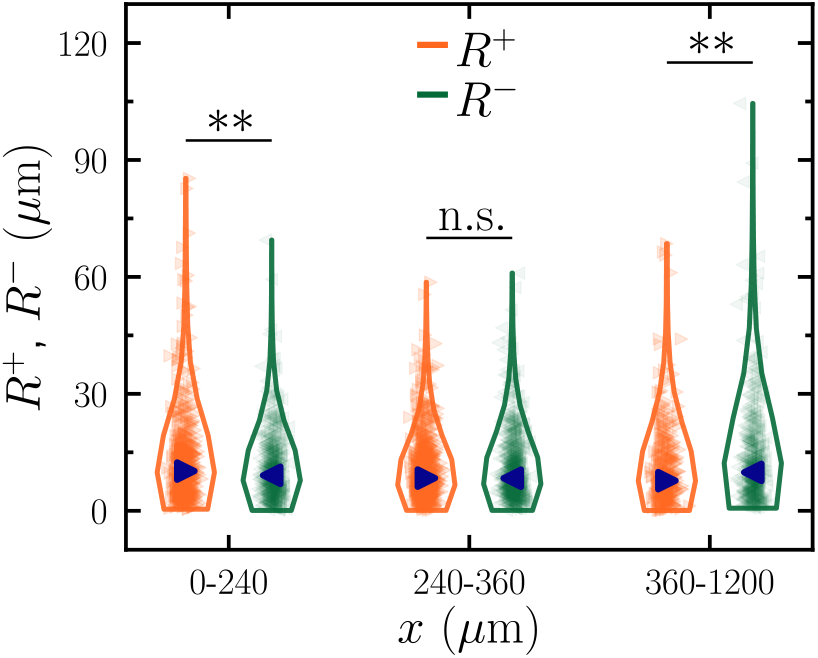
Distribution of run lengths for bacteria swimming to the right, away from the air-water interface (*R*^+^, orange right-pointing triangles), and to the left, toward the interface (*R*^−^, green left-pointing triangles), measured at three regions relative to the density peak: left (*x* = 0 − 240 *µ*m), middle (*x* = 240 −360 *µ*m), and right (*x* = 360 − 1200 *µ*m). Dark blue triangles indicate the median of each distribution. Bacterial density peaks at *x*_*p*_ ≈ 300 *µ*m. Statistical significance is assessed using a two-sided Kolmogorov-Smirnov test, where ** denotes *p <* 0.01 and ‘n.s.’ indicates no significant difference.

These results demonstrate that bacteria on either side of the density peak elongate their run lengths to swim toward the region of intermediate oxygen concentrations. Given *R*^+^ = *V*_0_*/λ*^+^ and *R*^−^ = *V*_0_*/λ*^−^, the observation of *R*^+^ *> R*^−^ on the left side of the peak suggests *λ*^+^ *< λ*^−^ and therefore a positive Δ*λ* = *λ*^−^ −*λ*^+^. Conversely, *λ*^−^ *< λ*^+^ on the right side of the peak, giving rise to a negative Δ*λ*. Figure 6 shows Δ*λ* estimated from the trajectories of the individual bacteria in the experiments, which clearly shows the sign change across the density peak. The magnitude of Δ*λ* is within the same order of magnitude as the density gradient (1*/V*_0_)*∂*_*x*_ log(*ρ*) extracted from experiments, in accordance with Eq. (18).

Our results indicate that the bacterial band is maintained by a mismatch in tumbling rates between left- and right-moving bacteria—a defining feature of aerotaxis that stands in contrast to the classical chemotactic mechanism captured by the Keller-Segel model [40, 66]. Crucially, the density gradient plays an active role in off-setting this tumbling imbalance, an effect not accounted for in earlier analyses [46]. These observations provide direct, single-cell-level evidence for the limitations of previous models (Sec. II B) and support the central assumption of optimal oxygen sensing in both the kinetic theory and the minimal model. More broadly, our results illustrate the importance of tracking individual bacteria in dense suspensions, revealing dynamics inaccessible to bulk measurements, and demonstrating the promise of this approach for future chemotaxis studies.

## VI. MOLECULAR MECHANISM FOR AEROTAXIS

In this last section, we explore the signaling pathway that enables optimal sensing and connect the understanding of signal processing at the molecular level, the motility of a single bacterium, and the spatiotemporal evolution of bacterial populations. We begin by identifying the potential mechanisms involved in the signaling pathway of *E. coli* that facilitate its aerotactic response, through the study of receptor-null mutants. Building on our findings, we then introduce a coarse-grained model of optimal sensing for aerotaxis, which predicts the steady-state position of the bacterial density peak and the sign change of Δ*λ* at the optimal oxygen concentration.

### A. Signaling pathway

*E. coli* adjusts the frequency of tumbling and extends runs in response to favorable chemical gradients. The transition between run and tumble is controlled by a signaling pathway that relays environmental cues to the molecular motors. Figure 9 illustrates the chemotaxis signaling pathway of *E. coli. E. coli* has five methyl-accepting chemotaxis proteins (MCPs): Tsr, Tar, Tap, Trg, and Aer. Each MCP is connected to the histidine kinase CheA via the linker protein CheW, forming a signaling complex. While the first four MCPs bind to external ligands, Aer does not have an external binding site [45] and likely regulates CheA activity via its methylation sites. CheA undergoes autophosphorylation to produce CheA-P, with its rate influenced by ligand binding. Attractant binding reduces CheA autophosphorylation, while repellent binding increases it, thereby transducing the external signals. CheA-P transfers its phosphate to the response regulator CheY, forming CheY-P. This phosphorylated form of CheY diffuses in the cell and binds to the motor, causing its switch from counter-clockwise to clockwise rotation, which triggers tumbling. The protein CheZ dephosphorylates CheY-P to CheY, preventing motor modulation. Thus, attractant binding reduces CheY-P levels, promoting smooth swimming to-ward favorable environments.

**FIG. 9.**
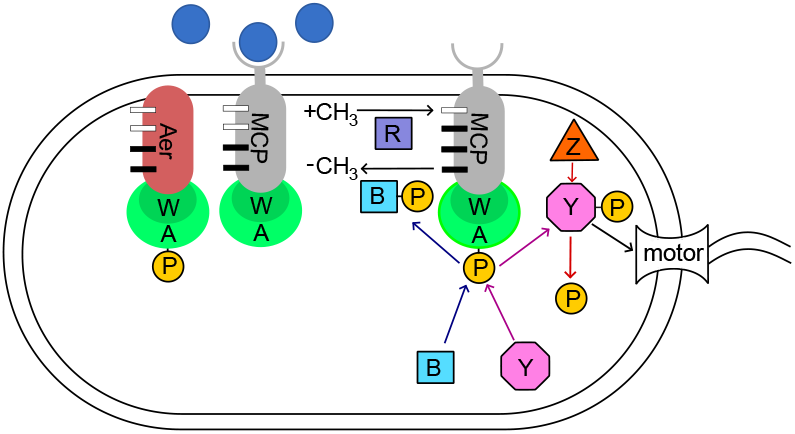
Schematic of the molecular signaling pathway of *E. coli* chemotaxis. Blue circles represent chemoattractant molecules in the environment. The labeled proteins—CheA (light green), CheW (dark green), CheR (dark blue), CheB (cyan), CheY (purple), and CheZ (dark orange)—are key components of the signaling pathway. P denotes a phosphate group. Methyl-accepting chemoreceptor proteins (MCPs) with external ligand-binding domains are depicted in gray, whereas the MCP Aer, which lacks an external ligand-binding domain, is shown in dark red. Methylation sites are represented by rectangular bars on the chemoreceptor, with filled and unfilled bars indicating occupied and unoccupied sites, respectively.

A sudden increase in attractant reduces tumbling frequency, followed by slow adaptation back to base-line. This adaptation is mediated by methylation and demethylation of MCPs, altering the kinase activity of CheA. CheR methylates the MCPs, while CheB-P removes methyl groups. When CheA autophosphorylation is high, CheB-P is formed, enabling negative feedback that counteracts excessive kinase activity. Thus, CheA kinase activity is regulated by (a) ligand binding and (b) methyl group attachment/detachment on MCPs.

The chemotaxis pathway described above can guide *E. coli* not only toward regions of high attractant or low repellent concentrations but also to locations with optimal intermediate levels of stimuli, such as pH [83] and temperature [84, 85]. Two mechanisms are known for optimal level sensing. In the first mechanism, two receptors with opposing responses to the same stimulus work together, allowing bacteria to seek an intermediate level of the stimulus. This mechanism operates in pH-taxis, where the Tar receptor directs movement toward low pH, while the Tsr receptor directs movement toward high pH [83]. In the second mechanism, a single receptor switches its response at a threshold stimulus level, therefore allowing bacteria to accumulate at the optimal stimulus level determined by this threshold. This mechanism underlies thermotaxis [86, 87].

Which mechanism governs *E. coli* aerotaxis? Oxygen sensing is mediated by the Aer and Tsr receptors [45]. To determine the mechanism, we measure the density profiles of bacteria lacking either the Aer receptor (Δ*aer*) or the Tsr receptor (Δ*tsr*). Figure 10 shows that even with only one functional receptor, bacteria still accumulate at an optimal oxygen concentration, suggesting that aerotaxis follows the second mechanism. Specifically, a single receptor (Aer or Tsr) inverts its response at a critical oxygen level, rather than the two receptors directing the movement of bacteria toward high and low oxygen levels, respectively.

**FIG. 10.**
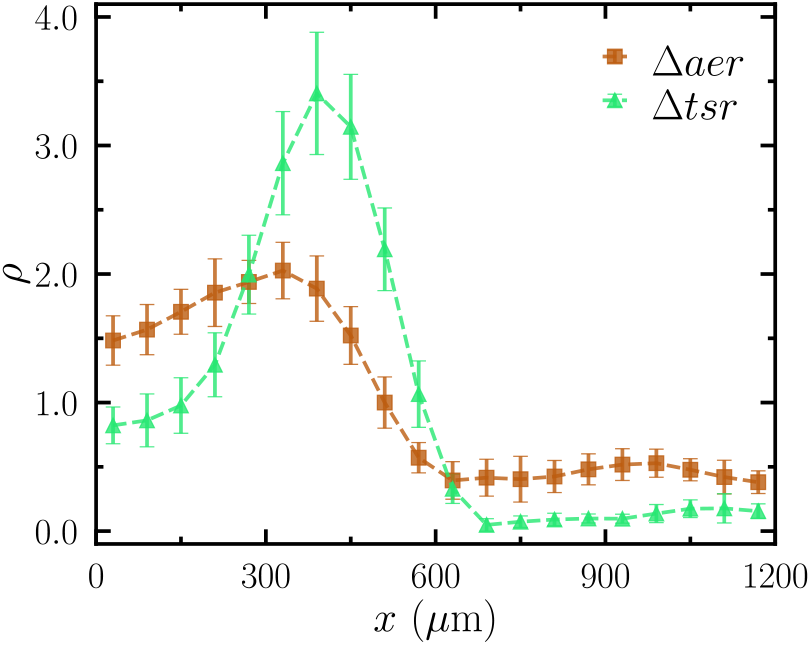
Steady-state density profiles of *E. coli* mutants lacking the Aer receptor (Δ*aer*, brown squares) and the Tsr receptor (Δ*tsr*, green triangles). The overall bacterial density is *ρ*_0_ = 4 × 10^9^ cells/ml, and the profiles are measured at *t* = 20 minutes. Error bars indicate the standard error.

### B. Molecular model

Guided by this experimental observation, we adopt the mathematical model initially developed for *E. coli* thermotaxis [86] to explain the accumulation of cells at an optimal oxygen concentration. Consider a 1D telegrapher’s equation describing bacterial populations, similar to that outlined in Sec. IV B. The present formulation extends the previous description by incorporating internal kinase activity as a new kinetic parameter. Here, the probability densities of cells traveling in the ± *x* direction at position ± *x*, and time *t* with internal kinase activity state *a* are denoted as *P* ^±^(*x, t, a*). Their evolution follows

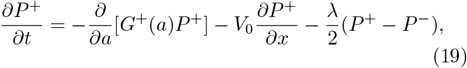

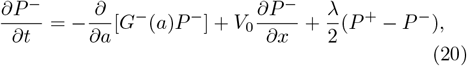

where *G*^±^ represents the time rate of change of kinase activity *a* for the population *P* ^±^, and *λ*(*a*) is the tumbling rate that in turn depends on the internal kinase activity. In this pathway-based description, the tumbling rate is a function of the kinase activity *a* that depends on the local oxygen concentration *c* externally and the methylation level internally. The local oxygen concentration sensed by a bacterium during a run, and thus the rate of change of *a*, is a Lagrangian measure that depends on the material derivative for each subpopulation *P* ^±^ [88]. Consequently, the differential Lagrangian measures give rise to different tumbling rates, *λ*^±^, for the two subpopulations, resulting in a coarse-grained description in Sec. IV B.

From Eqs. (19) and (20), Tu and co-workers [54, 89, 90] demonstrated that the population density of the bacteria obeys an effective advection-diffusion equation at long times, averaging over multiple tumble events

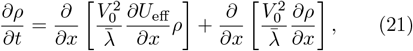

where *ρ*(*x, t*) = (*P* ^+^ + *P* ^−^)d*a* is the bacterial density, 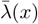 is the average tumbling rate in the population, and 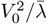 gives the effective diffusivity. More importantly, *U*_eff_ is an effective potential governing the advective flux, which depends on the tumbling frequency *λ* via the kinase activity *a* which in turn is regulated by the external oxygen concentration *c* and the internal methylation level, as described in Sec. VI A. The detailed derivation of the dependence of *U*_eff_ on the level of a stimulus (in this case, the concentration of oxygen) is intricate and has been previously presented in Refs. [86, 87]. To avoid redundancy, we summarize the key steps in the Supplemental Material (SM) and refer interested readers to the original publications for the full derivation.

Once *U*_eff_ is established from the signaling pathway, the steady-state density profile can be found from Eq. (21) by setting *∂ρ/∂t* = 0

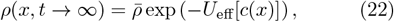

which is akin to a Boltzmann distribution with 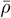 a normalization constant ensuring the conservation of the number of bacteria.

We evaluate the effective potential, *U*_eff_, from both the kinetic theory and the molecular model of the signaling pathway. Specifically, using the steady-state density profile for *ρ*_0_ = 1.5 × 10^9^ cells/ml at *t* = 20 minutes from the kinetic theory (Fig. 3(g)), we obtain *U*_eff_ directly by inverting Eq. (22), 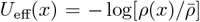. Alternatively, using the oxygen profile *c*(*x*) from the kinetic theory as input, we calculate *U*_eff_ through its dependence on the kinase activity in the molecular model (SM). Figure 11 compares the effective potentials derived from the two approaches. Since bacteria accumulate at the minima of these potentials, both approaches predict a similar location for the bacterial density peak. Extending these calculations to different overall densities *ρ*_0_, the molecular model successfully reproduces the position of the steady-state density peak at different *ρ*_0_, 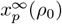, from the kinetic theory (Fig. 11 inset). The quantitative agreement establishes a strong correlation between molecular dynamics in the signaling pathway and bacterial accumulation at the optimal oxygen concentration on a population scale.

**FIG. 11.**
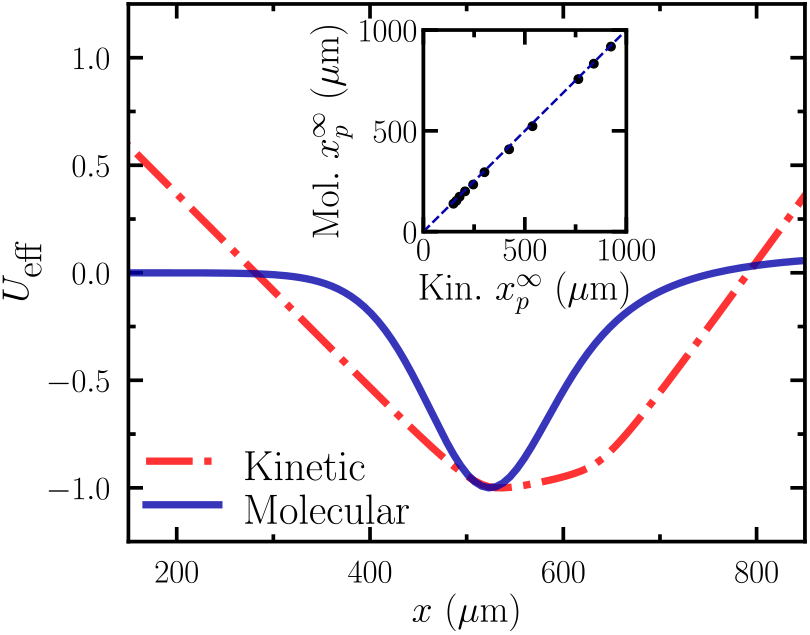
Effective potential, *U*_eff_, from the kinetic theory (the red dash-dot curve) and the molecular model (the blue solid curve). The potentials are normalized by their absolute minimum values. Inset: Steady-state positions of the bacterial density peak, 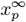, from the kinetic theory and the molecular model. The dashed line is the identity line.

Finally, comparing Eqs. (18) and (22) shows that the change of the tumbling frequency is related to the effective potential via Δ*λ* = −*V*_0_(*∂U*_eff_ */∂x*). The sign change of Δ*λ* corresponds to the minimum of *U*_eff_. Figure 6 presents Δ*λ* calculated from the molecular model alongside values obtained from experiments, the kinetic theory, and the minimal model. Notably, Δ*λ* changes sign at the bacterial density peak across all measurements and model calculations, highlighting the unification of the aerotaxis mechanism across different scales.

## VII. DISCUSSION

In this comprehensive study of bacterial aerotaxis, we demonstrate the accumulation of *E. coli* at an optimal oxygen concentration through experiments, theory, and simulations, linking molecular-scale signaling pathways, the run-and-tumble dynamics of individual bacteria, and population-scale collective migration. The sign change of the tumbling frequency difference Δ*λ* at an optimal oxygen concentration, coupled with the timescale separation between rapid oxygen diffusion and consumption and slow bacterial aerotactic response, drives the formation and migration of the bacterial band and ultimately leads to the reversal of migration direction as the overall bacterial density increases. By identifying the bacterial density as the key control parameter in this complex dynamic process, our work reconciles conflicting reports in the literature, which describe *E. coli* migrating toward both high [8, 40] and low [46] oxygen concentrations.

Our study highlights that bacterial migration direction and oxygen penetration are both influenced by bacterial density. Douarche *et al*. used oxygen-sensitive dyes to show that, at low bacterial densities, oxygen penetrates deeper into the channel [46], consistent with our model (Fig. 3(c)). It would be interesting to test whether oxygen retreats toward the air-water interface at high bacterial densities. Additionally, directly measuring oxygen concentration while tracking the run-and-tumble dynamics of individual bacteria would provide better estimates of the optimal oxygen concentration preferred by *E. coli*.

Our kinetic theory expands the optimal oxygen-seeking model, originally developed for the run-and-reversing bacterium *A. brasilense* [20], to the run-and-tumble dynamics of *E. coli*. It assumes a discontinuous variation in tumbling frequency across optimal oxygen levels (Eq. (4) and Fig. 3(b)). While the model quantitatively predicts the critical bacterial density, the oxygen-seeking behavior of *E. coli* is likely more complex. Future studies should aim to quantify oxygen-dependent run-and-tumble dynamics using a temporal assay [91], controlled oxygen release [92], or optical trapping to measure tumbling frequency within a spatially calibrated oxygen profile [93], as well as by probing intracellular oxygen levels with biosensors [94]. These measurements will inspire more precise models of bacterial aerotaxis.

Additionally, our model does not account for inter-bacterial steric and hydrodynamic interactions. At high densities, these interactions can induce flow instabilities [73, 74], leading to collective motion of bacteria known as bacterial turbulence [60, 95]. The threshold density for bacterial turbulence exceeds the peak density in our experiments [59], justifying the omission of inter-bacterial interactions in our model. However, even below this threshold, these interactions may still influence tumbling dynamics [96]. While our current model captures the trend of migration velocity (Fig. 2), incorporating these interactions could further improve its quantitative predictive capability.

We apply the optimal sensing model, initially developed for *E. coli* thermotaxis, to explore the molecular basis of aerotaxis. A key future direction is to investigate the signaling mechanisms underlying optimal oxygen seeking. FRET assays [97–99] could quantify molecular interactions in the aerotaxis signal transduction pathway, providing precise parameters for mathematical modeling. Moreover, future work could extend our 2D kinetic theory to incorporate the molecular sensing mechanism. Understanding the transient dynamics of this extended model would directly couple molecular-scale activity with population-scale collective motion. However, this poses challenges due to multiple timescales, from signaling dynamics on the millisecond scale to bacterial population dynamics on the minute scale. Our work provides a foundation for this development, with broader implications for multiscale theoretical understanding of sensing and population dynamics.

Finally, a critical bacterial density at which migration ceases could trigger a transition from a swimming state to a sessile state, fostering biofilm formation. Notably, the critical density, *ρ*_*c*_, around 1 × 10^9^ cells/ml observed in our experiments aligns with the native density of *E. coli* in the human gut [7]. This suggests that *ρ*_*c*_ may have been evolutionarily optimized to enhance the survival of *E. coli* in its native habitat, providing a potential physiological context for our findings.

## ACKNOWLEDGMENTS

We thank Gladys Alexandre, Vasilios Alexiades, Satyam Anand, Alex Mogilner, Yuhai Tu, and Tingtao Zhou for insightful discussions. D.G. and X.C. were supported by NSF CBET 1702352 and 2028652 and the David and Lucile Packard Foundation. B.C. acknowledges the support of the Department of Atomic Energy, Government of India, under project no. RTI4001.

## VIII. APPENDIX

### A. Cell culture

We prepared bacterial suspensions containing a mixture of fluorescently tagged and non-fluorescent wild-type *E. coli* strain AW HCB1732. The wild-type cells were tagged with green fluorescent protein (GFP) by inserting the plasmid pJUMP27-1A(sfGFP) (Addgene #126974). The bacterial suspensions were cultured by inoculating a small quantity of frozen stock in 2 ml Terrific Broth (TB) [tryptone 1.2% (w/v), yeast extract 2.4% (w/v), and glycerol 0.4% (w/v)]. The cell culture for the GFP-tagged cells was supplemented with 0.1% (v/v) of 50 mg/ml kanamycin. These cultures were then incubated in an orbital shaker at 250 rpm at 37^°^ for 12 to 16 hours in the late exponential phase. To ensure the motility of the cells, the bacterial suspensions were then diluted 1:100 with fresh TB and grown in the exponential phase for 6.5 h in the orbital shaker at 30^°^.

The mutant strains containing knockouts of the aerotaxis receptors, Δ*aer* and Δ*tsr*, were purchased from the *E. Coli* Genetic Stock Center and are part of the KEIO collection [100]. These strains were fluorescently-tagged using the plasmid pUCBB-eGFP (Addgene #32548). The culture media for the non-fluorescent cells of the mutant strains were supplemented with 0.1% (v/v) of 50 mg/ml kanamycin, whereas the fluorescent cells were cultured with 0.1% (v/v) each of 50 mg/ml kanamycin and 100 mg/ml ampicillin. The remainder of the culturing protocol is identical to that used for the wild-type cells.

At the end of the growth phases of cells, we mixed the liquid cultures containing the fluorescent and non-fluorescent strains of the bacteria. The volume of the two cultures for mixing was adjusted depending on the total concentration of cells. For each overall concentration, we ensured that the number of fluorescent cells in the field of view was optimized to have a sufficient number of cells for statistical averaging while allowing for the individual cell trajectories to be tracked without overlap from their neighbors. The motile cells were then harvested using gentle centrifugation at 800*g* for 5 min, followed by discarding the supernatant TB, and resuspending the cells in motility buffer [0.01 M potassium phosphate, 0.067 M NaCl, and 10^−4^ M EDTA, pH 7.0]. The suspension was then washed twice and the desired overall concentration of cells was obtained by adding motility buffer. The overall density of the bacterial suspension containing fluorescent and non-fluorescent cells in the motility buffer was measured using a biophotometer via optical density at 600 nm (OD_600_). We collected bacterial suspensions of different densities ranging from 6×10^8^ to 8×10^9^ cells/ml.

### B. Experimental setup and imaging

We used a glass capillary channel (Rectangle-Miniature Hollow Glass Tubing 5010, VitroTubes−) with a wall thickness of 0.07 mm, a vertical height of 100 *µ*m, and a horizontal width of 1.0 mm. The original channel is 50 mm in length, from which we cut a section with a length of 25 mm. Using UV-curable adhesive, we then attached this capillary tube to a microscope cover glass of 24 × 60 mm^2^. We filled the channel with a bacterial suspension and sealed one end with the adhesive, leaving the other end open for oxygen to diffuse into the channel. To minimize evaporation during the experiment, we covered the channel with an acrylic lid and placed soaked cotton wicks inside the chamber. The thin walls of the capillary channel allowed high-magnification imaging of individual bacteria.

The capillary channel filled with the bacterial suspension was mounted on a piezo stage with horizontal motion controlled using the Nikon NIS-Elements macro language. Bacteria in the channel were imaged using a Nikon Plan APO 60× oil objective (NA = 1.4) on an inverted Nikon Ti-Eclipse microscope with a Yokogawa CSU-X confocal spinning disk. The sample was excited using a 488-nm laser.

The field of view of our imaging is 160 × 120 *µ*m^2^. We wrote a custom macro script, allowing us to image swimming bacteria in the field of view for 10 seconds at 30 frames per second and then shift by 120 *µ*m to image an immediately adjacent region. We started imaging from the air-water interface at *x* = 0, and shifted the lens horizontally to image progressively deeper into the capillary channel. We imaged at 10 consecutive horizontal locations along the channel covering a total length of 1200 *µ*m. The entire scanning along the length of the channel, including the time taken to write video files on disk, took approximately 2 minutes. After waiting an additional 2 minutes, we scanned the channel again to obtain the spatial information at the next time instant. The combined duration of the channel scan and waiting time allowed data recording at 4-minute intervals, with timestamps at *t* = 4, 8, 12, 16, and 20 minutes. The duration of our observation was shorter than the division time of the bacteria, ensuring that the effects of growth and division can be neglected. We imaged the samples 25 to 30 *µ*m above the bottom wall of the capillary channel to avoid the influence of the boundary.

### C. Cell tracking and image analysis

The image stacks from confocal microscopy were processed using the Python library skimage. The GFP-tagged cells appeared as bright ellipsoids against a dark background (Fig. 1(b)). We binarized the video frames using the Niblack algorithm [101] and obtained the centroid positions and orientations of the cells. The centroids of individual bacteria across consecutive frames were linked using a standard particle tracking algorithm to yield their swimming trajectories. We obtained the swimming speed, *V*_0_, and the swimming direction, *θ*, using the central finite difference between bacterial positions, and the arccosine of the orientation of bacterial displacement relative to the *x* axis of the lab frame.

### D. Bacterial density profile

For a given timestep of a scan along the channel, we recorded the spatial coordinate of each fluorescently tagged bacterium detected over the scanning duration. The total length of the observed channel of 1200 *µ*m was binned into 60 *µ*m windows and the number of bacteria at each of the *n*_*w*_ = 1200*/*60 = 20 windows, *n*(*x*_*i*_), was calculated. To reduce noise from fluctuations in the observed bacterial count, we applied a Savitzky-Golay filter, fitting a 5th-order polynomial with a window size of 9, which yielded the spatially smoothed bacterial count, *n*_*s*_(*x*). Finally, the normalized local density of bacteria was calculated as *ρ*(*x*) = *n*_*s*_(*x*)*/*(*N*_*s*_*/n*_*w*_), where 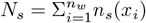 is the total number of bacteria observed along the length of the channel obtained after spatial smoothing. The normalized density eliminated the influence of the variation of the ratio of the fluorescent to non-fluorescent bacteria for different overall bacterial densities.

### E. Migration speed of bacterial population

To obtain the population migration speed in experiments, *v*_*m*_, we first calculated the mean position of the bacterial population 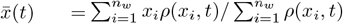. Finally, *v*_*m*_ was obtained through a linear fit of 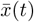 versus *t*. The uncertainty in the slope of the linear fit was taken as the experimental error for *v*_*m*_.

Similarly, to evaluate the population migration speed from the kinetic theory, we first obtained the mean position of bacterial population using 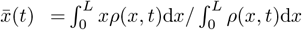. The instantaneous speed was then obtained as 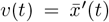. The average population migration speed was calculated following 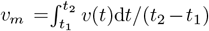 over a time interval [*t*_*1*_, *t*_*2*_]. The lower time limit *t*_1_ was chosen to exclude the influence of the transient dynamics at short times. For *ρ < ρ*_*c*_, the bacterial density peak shows unidirectional migration away from the interface (Fig. 4a). In this low-density regime, we chose the inflection point of 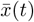 as *t*_1_, i.e. 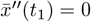. For *ρ > ρ*_*c*_, the peak of the density profile *x*_*p*_ is non-monotonic at early times. Here, we chose *t*_1_ as the time where *x*_*p*_ reaches its maximum, i.e. 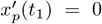 and 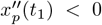. For both cases, the upper time limit *t*_2_ was chosen such that 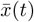 has reached steady state with *v*(*t > t*_2_) *<* 10^−4^ *µ*m/s.

### F. Δ*λ* from the kinetic theory

For the kinetic theory, we considered two subpopulations of bacteria, one consisting of bacteria swimming in the +*x* direction away from the air-water interface, and another consisting of bacteria swimming in the −*x* direction toward the interface. We then calculated the tumbling rates of the bacteria swimming along the +*x* and −*x* directions as 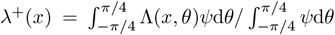 and 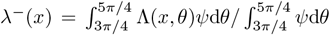, respectively. Here, *ψ*(*x, θ*) is the probability density function. Finally, we computed the difference in the tumbling rates between the two subpopulations at each spatial location,

Δ*λ*(*x*) = *λ*^−^(*x*) − *λ*^+^(*x*).

### G. Varying *α* and *γ* in the minimal model

We examined how the critical bacterial density *ρ*_*c*_ depends on the parameters *α* and *γ* in the expression Δ*λ* = *α* tanh[*γ*(*c − c*^*^)] in our minimal model. Here, *α* sets the magnitude of Δ*λ*, while *γ* controls the steepness of the variation of Δ*λ* near *c* = *c*^*^. We focused on the regime where the crossover between the initial peak position, 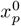, and the final peak position, 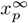, occurs within the experimental domain of 0 *< x <* 1200 *µ*m. We found that varying 0.02 *< α <* 0.06 s^−1^ and 0.05 *< γ <* 5.55 *µ*M^−1^ leads to *ρ*_*c*_ generally agreeing with our experimental observation (Table 1).

**TABLE 1.**
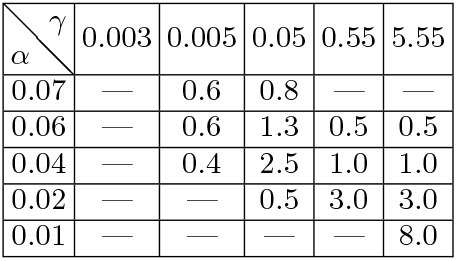
*ρ*_*c*_ (in 10^9^ cells/ml) as a function of *α* (s^−1^) and *γ* (*µ*M^−1^). ‘—’ indicates that the migration direction of the density peak does not reverse in the 1200 *µ*m-long channel.

| $\alpha \backslash \gamma$ | 0.003 | 0.005 | 0.05 | 0.55 | 5.55 |
| --- | --- | --- | --- | --- | --- |
| 0.07 | — | 0.6 | 0.8 | — | — |
| 0.06 | — | 0.6 | 1.3 | 0.5 | 0.5 |
| 0.04 | — | 0.4 | 2.5 | 1.0 | 1.0 |
| 0.02 | — | — | 0.5 | 3.0 | 3.0 |
| 0.01 | — | — | — | — | 8.0 |

## Notes

### Competing Interest Statement

The authors have declared no competing interest.

