## Supplemental Material for "Breathe in, breathe out: Bacterial density determines collective migration in aerotaxis"



(Dated: April 2, 2025)


### THE MOLECULAR MODEL OF SIGNALING PATHWAY

We summarize below the key steps in the derivation of the effective potential $U_{\text{eff}}$ and its dependence on the oxygen concentration $c(x)$ from the signaling pathway. We refer interested readers to the original publications [1, 2] for the full derivation.

Particularly, we extend the original derivative of $U_{\text{eff}}$ for thermotaxis [1, 2] to aerotaxis, where the spatial gradient of $U_{\text{eff}}$ is given as

$$\frac{\partial U_{\text{eff}}}{\partial x} = \frac{1}{\bar{\lambda}} \left( \frac{\partial \lambda}{\partial a} \frac{\partial a}{\partial c} \right)_{a=\bar{a}} \frac{\partial c}{\partial x}. \tag{1}$$

Here, $\bar{\lambda}$ is the average tumbling rate of bacteria. The derivatives are evaluated at the average kinase activity $\bar{a}$. To calculate $U_{\text{eff}}$, we need to consider the biochemical factors in the signaling pathway described in Sec. VIA of the main text to relate the tumble frequency $\lambda$ to the kinase activity $a$ and to determine the dependence of $a$ on the oxygen concentration $c$. Table S1 lists the key parameters used in the molecular model.

TABLE S1. Parameters related to the kinase activity, $a$, methylation level, $m$, and oxygen concentration, $c$, used in the molecular model.

| Symbol | Parameter |
|---|---|
| $a$ | Instantaneous kinase activity |
| $\bar{a}$ | Steady-state average kinase acitivity |
| $m$ | Instantaneous methylation level |
| $\bar{m}$ | Steady-state average methylation level |
| $m^*$ | Critical methylation level for response inversion |
| $\hat{m}$ | Reference methylation level for $f_a(\hat{m}, \hat{c}) = 0$ |
| $c$ | Oxygen concentration |
| $c^*$ | Critical oxygen concentration for response inversion |
| $\hat{c}$ | Reference oxygen concentration for $f_a(\hat{m}, \hat{c}) = 0$ |

First, the tumbling frequency $\lambda$ as a function of the kinase activity $a$ is given as,

$$\lambda = d_r + \frac{1}{\tau_\lambda} \left( \frac{a}{K_{1/2}} \right)^H, \tag{2}$$

where $\tau_\lambda$ is the duration of a tumble, $K_{1/2}$ is the activity level for a motor with a probability of clockwise rotation of 0.5, and $H$ is the Hill coefficient for the response of the motor to kinase [3].

Second, the kinase activity $a$ depends on the external oxygen concentration, $c$, as well as the internal methylation level, $m$, through the free energy $f_a(m, c)$ of the chemotactic receptors as,

$$a(m, c) = \frac{1}{1 + \exp[N f_a(m, c)]}, \tag{3}$$

where $N$ is the number of dimers of chemoreceptor proteins constituting a single receptor unit. To accumulate at an optimal intermediate oxygen concentration at steady state, the oxygen dependence of the kinase activity must undergo an inversion at a critical oxygen concentration $c = c^*$, which is associated with a critical methylation level $m = m^*$ at steady state (see Eq. (8) below). The simplest mathematical form for $f_a(m, c)$ that allows this inversion of response is a linear expansion about a reference methylation level $\hat{m}$ and reference oxygen concentration $\hat{c}$. Thus, the free energy for an individual receptor that responds to oxygen (e.g. Aer or Tsr) can be approximated as:

$$f_a(m, c) \approx -E_m(m - \hat{m}) - \alpha_0(m - m^*)(c - \hat{c}), \tag{4}$$

where $\alpha_0$ and $E_m$ are positive constants. The reference values $\hat{m}$ and $\hat{c}$ determine $f_a(\hat{m}, \hat{c}) = 0$. Thus, we have

$$\frac{\partial a}{\partial c} = -Na(1 - a) \frac{\partial f_a}{\partial c} = N\alpha_0 a(1 - a)(m - m^*). \tag{5}$$

The dynamics of methylation and demethylation are much slower compared to the average run time of an individual bacterium [4, 5]. Following Ref. [6], this separation of time scales enables us to calculate an average steady-state methylation level, $\bar{m}(c)$, which depends only on the local oxygen concentration, $c$. Substituting $\bar{m}(c)$ in Eq. (3) then yields the average steady-state activity level, $\bar{a}(c) \equiv a(\bar{m}, c)$.

Finally, to obtain $U_{\text{eff}}$, we need to evaluate $\partial a/\partial c$ at the average steady-state kinase activity $\bar{a}(c)$. As described in Sec. VI A of the main text, methylation regulates the kinase activity $a$ close to its adapted value $a_0$, which can be described mathematically as $\mathrm{d}m/\mathrm{d}t = (a_0 - a)/\tau_m$, where $\tau_m$ is the adaptation time. The steady-state methylation level is obtained from $[a_0(c) - a(\bar{m}, c)]/\tau_m = 0$ or equivalently $\bar{a}(c) = a_0(c)$. We assume that the adapted activity depends on oxygen concentration as

$$a_0(c) = \frac{1}{1 + \exp[-\sigma(c - \hat{c})]}, \tag{6}$$

where $\sigma$ is a constant. Equation (6), originally derived for thermotaxis, predicts that at low oxygen concentrations, $a_0(c)$ would decrease and the bacterium would tumble less frequently. Indeed, low oxygen concentrations reduce the proton motive force driving the *E. coli* flagellar motor [7]. Experiments have shown that the tumbling frequency is indeed reduced at low values of proton motive force [8], and, consequently, at low oxygen concentrations.

Combining Eqs. (3) and (6), we have $Nf_a(\bar{m}, c) = -\sigma(c - \hat{c})$, yielding,

$$\bar{m}(c) = \frac{(N\alpha_0 m^* + \sigma)(c - \hat{c}) + NE_m\hat{m}}{N[\alpha_0(c - \hat{c}) + E_m]}. \quad (7)$$

The average kinase activity, $\bar{a}$, can be computed by substituting $m = \bar{m}$ in Eqs. (3) and (4). The average tumbling frequency, $\bar{\lambda} = \lambda(\bar{a})$, is then obtained from Eq. (2). For a given $c(x)$, $U_{\text{eff}}(x)$ can finally be calculated from Eq. (1) using Eqs. (2) and (5).

Note that the steady-state methylation at the critical oxygen concentration gives the critical methylation level, $\bar{m}(c^*) = m^*$. Thus, from Eq. (7), $c^*$ and $m^*$ are related by,

$$c^* = \hat{c} + NE_m(m^* - \hat{m})/\sigma. \quad (8)$$

From Eq. (5), we see that at steady state, $\frac{\partial a(\bar{m},c)}{\partial c} > 0$ for $\bar{m} > m^*$ and $\frac{\partial a(\bar{m},c)}{\partial c} < 0$ for $\bar{m} < m^*$, i.e. the inversion of the response of the kinase activity to oxygen at $m^*$ and therefore at $c^*$, necessary for the formation of a bacterial density peak at $c^*$.

To compare the prediction from the molecular signaling pathway with the results from the kinetic theory, the following parameters on the methylation and run-and-tumble dynamics of *E. coli* are taken from Ref. [1], namely, $\hat{m} = 1.94$, $E_m = 1$, $m^* = 2.2$, $N = 6$, $d_r = 0.28$ $s^{-1}$, $\tau_\lambda = 0.2$ s, and $H = 10$. Specific to the oxygen response, we choose the parameters $\hat{c} = 0.02$, $\alpha_0 = 40$, and $\sigma = 20$.
